# Effects on brain activity after creative mathematical reasoning when considering individual differences in cognitive ability

**DOI:** 10.1101/2021.03.30.437492

**Authors:** Linnea Karlsson Wirebring, Carola Wiklund-Hörnqvist, Sara Stillesjö, Carina Granberg, Johan Lithner, Micael Andersson, Lars Nyberg, Bert Jonsson

## Abstract

Many learning opportunities of mathematical reasoning in school encourage imitative learning procedures (*algorithmic reasoning, AR*) instead of engaging in more constructive reasoning processes (e.g., *creative mathematical reasoning, CMR*). Here, we employed a within-subject intervention in the classroom with pupils in upper secondary schools followed by a test situation during brain imaging with fMRI one week later. We hypothesized that learning with CMR compared to AR should lead to a CMR-effect, characterized by better performance and differential brain activity during test. We observed higher brain activity in key regions for mathematical cognition such as left angular gyrus and left inferior frontal gyrus on tasks previously learnt with CMR compared to AR. The effects remained when controlling for individual differences in cognitive abilities, as well as performance and response time differences between the two conditions. Encouraging pupils to engage in constructive processes when learning mathematical reasoning might thus have lasting beneficial effects.

## 1. Introduction

Mathematics is a core subject in most countries’ educational systems. Still, when it comes to learning and instruction of mathematics in school, and how tasks in mathematics textbooks are commonly constructed, arguably more could be done to promote active learning (Freeman et al., 2014). It has been observed that many learning opportunities within the subject of mathematics encourage rote-learning or imitative, algorithmic procedures based on following solution templates presented and quite often explained by teachers and/or textbooks (Boesen et al., 2014; Hiebert, 2003; Jäder, Lithner, & Sidenvall, 2019). The term ‘algorithm’, which can have different meanings in different areas, here denotes a finite sequence of executable instructions for solving a task. As an example, Jäder et al. (2019) analyzed mathematical textbooks from 12 countries and concluded that nearly 80% of the tasks promoted imitating a provided algorithmic solution template rather than actively engaging the pupil in more constructive reasoning processes. But how could such constructive learning of mathematical reasoning be promoted, and would it have effects on learning and memory?

A way of characterizing a constructive form of mathematical reasoning for solving training tasks, and corresponding ways to design teaching and textbook tasks, has been suggested and developed within the *Learning by Imitative and Creative Reasoning research program* (*LICR*; Lithner, 2008; 2017). The main focus within the LICR-program is to challenge the dominant way of learning mathematics through imitative learning strategies, with tasks promoting *creative mathematical reasoning* (CMR-tasks; see e.g., Lithner, 2008; 2017). In contrast to tasks promoting imitating a provided algorithmic solution template, CMR-tasks are designed to favor active learning because a solution template is *not* given to the pupil as a part of the task. Instead, the pupils are forced to novel mathematically based reasoning by actively constructing the solution method themselves. Eye-tracking studies report on significant differences comparing students’ approach to solving AR tasks compared to CMR tasks. In AR tasks students tend to choose algorithmic reasoning following the provided template even though creative reasoning is an available choice. Students solving AR tasks only pay attention to the template, they do not, compared to students solving CMR tasks, focus on the task-relevant information they would need in order to create a method and reach a deeper understanding of the solution (e.g. Norqvist, Jonsson, Lithner, Qwillbard & Holm, 2019). Empirical evidence have confirmed that CMR-tasks, compared to the more passive mode exemplified above: *algorithmic reasoning* (AR, following a given or recalled algorithm), have proven beneficial for mathematical learning (e.g., Jonsson, Granberg, & Lithner, 2020a; Jonsson, Kulaksiz, & Lithner, 2016; Jonsson, Norqvist, Liljekvist, & Lithner, 2014; Norqvist, 2018; Norqvist et al., 2019; Wirebring, Lithner, et al., 2015). Similar results have been reported even when the AR tasks were followed by explanations of the mathematics supporting the suggested templates (Norqvist, 2018).

The importance of evidence-based teaching strategies that are also plausible from a neurophysiological perspective has been highlighted with the emerging field of *educational neuroscience* (e.g. Thomas et al. 2019). With brain imaging data, it becomes possible to observe neural evidence for differences in learning outcomes despite relatively small behavioral differences, especially in relation to the complexity of mathematical thinking (Sohn et al., 2004). Brain imaging support for a difference in how the brain is engaged during mathematical reasoning after initial training with CMR-tasks versus AR-tasks was obtained in a previous study (Wirebring, Lithner, et al., 2015). A fronto-parietal brain network was commonly engaged when participants solved mathematical test questions, which is consistent with other findings that the inferior frontal gyrus (IFG) and angular gyrus (AG) are recruited during advanced mathematics (Zhou et al., 2018). In addition, differences in recruitment of the left AG were observed as a function of learning method (AR > CMR). The left AG has repeatedly been implicated in studies of simple arithmetical learning (see e.g. Zamarian, Ischebeck, & Delazer, 2009). Differential AG activity in the Wirebring, Lithner, et al. (2015) study was interpreted in relation to the notion that activity in this region is modulated by the demands on verbal retrieval strategies (Dehaene, Piazza, Pinel, & Cohen, 2003). Higher activity in left AG has been observed for practiced compared to novel arithmetic problems (Delazer et al., 2003; Ischebeck et al., 2006). Left lateralized AG activity has been seen to mirror memory retrieval-based strategies rather than computational strategies in multiplication (Grabner et al., 2009) and increases in left AG activity has been observed as a function of learning multiplication problems (Ischebeck, Zamarian, Egger, Schocke, & Delazer, 2007). AG activity after initial learning of mathematics through CMR tasks might be related to mobilization of relevant control networks (Kim, 2020), allocation of attention to memory (Cabeza, Ciaramelli, Olson, & Moscovitch, 2008) as well as reactivation of integrated mathematical representations related to terminologies and principles (Zhou et al., 2018). Seghier (2013) has proposed a unified framework for the multiple functions ascribed to the AG.

Interestingly, a relation between individual differences in cognitive abilities and brain activation in parietal and frontal cortices during mathematical processing has been observed (Grabner et al., 2007; Grabner, Reishofer, Koschutnig, & Ebner, 2011; Gullick, Sprute, & Temple, 2011; Price, Mazzocco, & Ansari, 2013; Wirebring, Lithner, et al., 2015). For example, Grabner et al. (2007) identified a positive relation between mathematical and numerical IQ and signal change in AG during arithmetic problem solving. Thus, the way the brain is activated during mathematical reasoning might be modulated by individual differences in cognitive ability.

Here we present a large-scale study of brain activity during mathematical reasoning one week after initial CMR or AR training. Because an interaction between cognitive abilities and learning methods that require more cognitive effort (e.g., CMR) cannot be ruled out, we opted for a factorial design (mathematical learning method by cognitive ability). Based on the relevant evidence from previous studies, we hypothesized differential recruitment of the left AG region. A more open question was the direction of the effect and the potential for interaction with cognitive ability.

## 2. Materials and Methods

### 2.1 Participants

The current study is part of a large data collection aimed at answering questions related to cognitive learning strategies (see also Jonsson et al., 2020a; Jonsson, Wiklund-Hörnqvist, Stenlund, Andersson, & Nyberg, 2020b; Wiklund-Hornqvist, Stillesjö, Andersson, Jonsson, & Nyberg, 2021). At the outset, 324 pupils were recruited for the data collection from third year classes in local upper secondary schools. The pupils were enrolled in programs with a focus on the natural sciences, technology and economics. Out of these participants, a subsample of 72 participants (32 male; *M*_Age_=18.0, *SD* = .41) took part in the mathematical learning intervention (Day 1), volunteered for fMRI (Day 7), and met the inclusion criteria for fMRI, i.e., all participants were neurologically healthy, right-handed by self-report and had normal or corrected-to-normal vision. Two participants had incomplete data and could not be included in analyses on cognitive ability (see below under *Cognitive Ability Measures*). fMRI data from six participants had to be corrected for motion with the ArtRepair algorithm (Mazaika, Whitfeld & Cooper, 2005). Data from one participant had to be discarded from the fMRI data analyses due to extensive head movements. All participants signed a written informed consent before participation. For participants who had not attained a legal age of majority (18 years; *n* = 5), written informed consent was obtained from the participant and both caregivers. The study was conducted in accordance with the Declaration of Helsinki and approved by the Regional Ethical Review Board in Umeå (ref number, 2016/259-31).

### 2.2 Design, Procedure and Materials

Prior to the learning intervention, all 324 participants were first tested with an extensive battery of cognitive ability measures (see Fig. 1a, and below for details). For the learning intervention, the study entailed a within-subject design where each participant learned to solve mathematical tasks in two learning conditions: AR and CMR (see Fig. 1a and Supplementary Fig. S1). At Day 1, participants learned to solve mathematical tasks in the classroom, and one week later, participants came back to be tested on the same tasks while undergoing fMRI (see Fig. 1b for the fMRI trial design).

**Figure 1.**
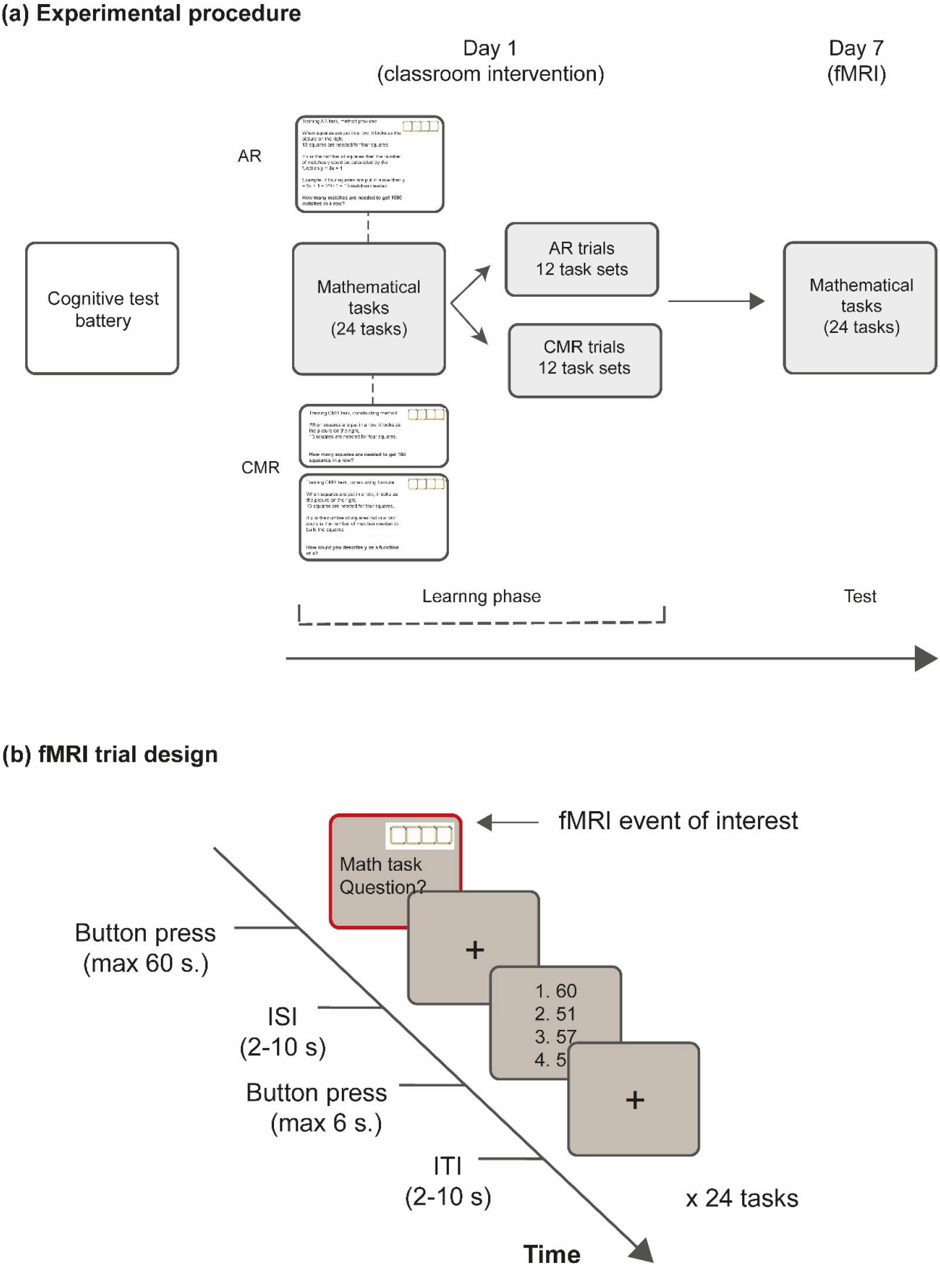
Experimental setup. a) Experimental procedure, b) fMRI trial design. AR = algorithmic reasoning. CMR = creative mathematical reasoning. ISI = interstimulus interval. ITI = intertrial interval. For larger versions of the task examples, see Supplementary Fig. S1. For additional examples, see Fig. S2.

#### 2.2.1 Cognitive Ability Measures

In order to capture cognitive abilities in a broad sense, nine different cognitive ability measures were administered as computerized tests a few weeks prior to the learning intervention. These included measures of *general fluid intelligence* (Ravens APM: Raven, Raven, & Court, 1998), *working memory capacity* (Operation span: Unsworth, Heitz, Schrock, & Engle, 2005), *episodic memory* (Modified associative learning test: Lövdén, 2003), *processing speed* (Number- and letter comparison: Schmiedek, Lövdén & Lindenberger, 2010), *visuospatial short-term memory* (Block span: Nyroos, Jonsson, Korhonen, & Eklöf, 2015; Wiklund-Hörnqvist, Jonsson, Korhonen, Eklöf, & Nyroos, 2016), *verbal short-term memory* (Digit span: Nyroos et al., 2015; Wiklund-Hörnqvist et al., 2016) and *updating* (Letter running span: Dahlin, Neely, Larsson, Backman, & Nyberg, 2008; Keep track: Miyake et al., 2000). For a description of each of these tasks, and the implementation and procedure, see Jonsson et al. (2020b).

To be able to relate individual differences in cognitive ability, in a broad sense, to the effects of the mathematical learning intervention on later performance and brain activity, we relied on a composite score based on performance on the cognitive ability measures mentioned above. This composite score was the individual factor score from a principal component analysis (PCA) run on the complete sample (*N* = 324; Jonsson et al., 2020b). The larger sample was divided into three equal groups based on each participant’s value on the factor score, denoted *low*, *intermediate* and *high* cognitive ability groups. Two participants had incomplete data and could not be categorized. Importantly, each remaining participant in the current fMRI study (*N* = 70) was categorized as low (*n* = 17), intermediate (*n* = 21) or high (*n* = 32) relative to the larger sample, making the categorization of participants in the current study less biased and more robust. In addition, a rationale for categorizing the participants in the current study into the three groups was to be able to relate to the study by Jonsson et al. (2020b) also theoretically. Jonsson and colleagues observed that the magnitude of the *testing effect* (the effect of another active learning method which compares vocabulary learning using retrieval practice with study) was independent of cognitive ability (low/intermediate/high), and we wanted to be able to investigate whether similar conclusions would follow when comparing CMR to AR (i.e. the CMR-effect). For details concerning distribution and intercorrelations of the cognitive tests, and details of the PCA, see Jonsson et al. (2020b).

#### 2.2.2 Mathematical Training

Each participant was presented to 24 mathematical tasks in two learning conditions: the AR condition (12 tasks) and the CMR condition (12 tasks) (for task illustration, see Fig. 1a and Supplementary Fig. S1 and S2). Participants practiced individually in the classroom in front of a laptop, with school desks spaced apart. The matching of tasks with learning condition was counterbalanced, such that half of the participants were presented to the tasks the other way around (i.e., the 12 tasks used for the AR condition were used for CMR and vice versa). The counterbalance was successful such that about half of the participants got certain 12 tasks for CMR and the other half received these tasks for AR, and vice versa. All participants started practicing in the AR condition, and after a short break continued with the CMR condition. The order was set like this in order to prevent carry-over effects from CMR to AR. This was motivated by the fact that most pupils should be acquainted with the elements of AR, because this is a common way of learning mathematics (e.g., Jäder et al., 2019), while CMR represents the more novel task design (Lithner, 2008; 2017). The set-up and the length of the intervention (e.g., the number of tasks to solve) was designed to be suitable as an in-class intervention and to avoid fatigue (the intervention lasted approximately 50 minutes (AR) + short break + 50 minutes (CMR)).

The general design elements of the AR and CMR conditions have been described in detail in several previous papers (see e.g., Jonsson et al., 2020a; Jonsson et al., 2016; Jonsson et al., 2014; Lithner, 2017; Norqvist, 2018; Norqvist et al., 2019; Wirebring, Lithner, et al., 2015). The key difference between AR and CMR is that while the mathematical tasks in the AR condition provide a solution template, that is, a formula and an example, this part is completely left out in the CMR condition of the task. In the CMR condition, the participant is instead asked to create their own solution method and furthermore asked to construct the corresponding formula (see Fig. S1 and Fig. S2).

Each task consisted of several sub-tasks, where the layout was identical but where the exact numbers that should be calculated differed. The first sub-task in both conditions was always numerically easy, but the following sub-tasks asked for a numerical solution where it was important to have a solution method at hand, as the number grew too large to come up with a correct answer by, for example, imagining and counting the matches needed to construct 100 squares (see Fig. 1a; S1; S2). The third sub-task of each task in the CMR condition always asked the participant to construct their own solution formula (Fig. S1b). Independent of learning condition (CMR or AR), participants were given four minutes to learn the various sub-tasks of each of the 24 tasks. If the pupils managed to solve all the sub-tasks, the software re-sampled sub-tasks until the 4 minutes had passed. Thus, an equal time-on-task for both learning conditions was ensured for all participants. After each answer given, participants received feedback on the correct answer. However, no correct answers were provided to tasks that asked the pupils to construct formulas (i.e., the third CMR task, the formula question; see Fig S1b). All participants had access to a small calculator on the side of the screen (during training but not during fMRI test) in order to even out differences in mental calculation skills as a hinder for learning and also to more closely mimic a real school situation.

#### 2.2.3 Mathematical Testing with fMRI

One week after the mathematical learning intervention, the participants came to the fMRI scanner to undergo fMRI while solving randomly presented test instances of the 24 tasks they had previously trained on Day 1 (Fig. 1b). The test instances did not provide a formula or an example, and participants had to recreate the solution method themselves. Participants solved one test instance per task with 60 seconds at their disposal (see Fig. 1b). The exact numbers used in the test tasks were different from those used during training. Participants were instructed to push a button on a response pad when they had solved the task and, after a variable delay, they were shown four numerical response alternatives. Participants marked which of the alternatives was the correct one and, after a variable delay, moved on to the next task (Fig. 1b).

Upon completion of the fMRI protocol and the structural images, participants filled out a follow-up questionnaire outside the scanner. For each of the 24 task sets, participants were asked how difficult they thought it was on a scale from 1 (very easy) to 7 (very difficult).

### 2.3 fMRI Image acquisition

FMRI image acquisition was made as in Wiklund-Hörnqvist et al. (2021). Image acquisition was made on a 3T GE Discovery MR 750 scanner (General Electrics). Functional T2*-weighted images were obtained with a single-shot gradient echo EPI sequence used for blood oxygen level dependent imaging. The following parameters were used for the sequence: echo time: 30 ms, repetition time: 2000 ms (37 slices acquired), flip angle: 90°, field of view: 25 x 25 cm, 96 x 96 matrix and 3.4 mm slice thickness. A 32 channel SENSE head coil was used. Signals arising from progressive saturation were eliminated through ten “dummy scans” performed prior to the image acquisition. The stimuli were presented on a computer screen that the participants viewed through a tilted mirror attached to the head coil. Presentation and reaction time data were handled by a PC running E-Prime 2.0 (Psychology Software Tools, Inc., USA) and fMRI optical response keypads (Current Designs, Inc., USA) were used to collect responses.

T1-weighted images were obtained with a 3D fast spoiled gradient echo sequence (FSPGR) in axial orientation. The following parameters were used: echo time: 3.2 ms, repetition time: 8.2 ms (176 slices acquired), flip angle: 12°, field of view: 25 x 25 cm, 256 x 256 matrix and 1 mm slice thickness.

### 2.4 fMRI Data analyses

The data was analyzed using SPM12 (Wellcome Department of Cognitive Neurology, UK) implemented in Matlab (R2014b) (Mathworks Inc., USA) and run through an inhouse program (DataZ). All images were corrected for slice timing, realigned to the first image volume in the series, and unwarped. Images were spatially normalized with Dartel (Ashburner, 2007). The individual T1 images were segmented, and a group specific mean template and individual flow fields were created with the DARTEL algorithm (Ashburner, 2007). The DARTEL template was used to normalize images to the anatomical space defined by the MNI atlas (SPM12), and smoothed using an 8.0 mm FWHM Gaussian filter kernel. Data were high-pass filtered with a cut-off of 128 s.

The general linear model consisted of two effects of interest (tasks learnt in the AR condition and tasks learnt in the CMR condition, Fig. 1b) and six effects of no interest (AR ISI, AR alternative forced-choice, AR ITI, CMR ISI, CMR alternative forced-choice, CMR ITI). We chose to model the blocks associated with the two conditions as separate effects in order to incorporate potential differences during the ITI, alternative forced-choice and ISI that would not be of relevance to the core question addressed. Six movement parameters were included as covariates of no interest. All regressors, except the movement parameters, were convolved with a hemodynamic response function. The time points when the mathematical task was shown were used as onset, and response times were used as duration. We used these variable durations in order to allow the analyses to capture the complete reasoning process, which could vary somewhat in duration from trial to trial and between participants.

In the first level analysis, model estimations were made for each participant. The individual model estimations were then taken into a second-level mixed model ANOVA with learning condition (AR/CMR) as within subject factor and cognitive ability group (low/intermediate/high) as between subject factor (unequal variances assumed between the groups). With this analysis, it is possible to investigate whether there is a main effect of learning condition (AR/CMR) on the brain activity data acquired one week after learning. Moreover, it also enabled us to examine whether there is main effect of cognitive ability group and possible interactions between learning condition and cognitive ability group.

The statistical threshold was set to *p* < .01 (FDR-corrected) at the voxel level, and *k* > 15 at the cluster level. The analysis was explicitly masked with a grey matter mask derived from a segmentation in SPM of the average of the individual MNI-normalized anatomical images of the participants in the study.

#### 2.4.1 Control Analyses

For completeness, we related the composite score of cognitive abilities to performance and brain activity by including the score as a continuous variable. In terms of performance, we ran a regression analysis to investigate to what extent the cognitive composite score predicted the difference in performance between CMR and AR (the *CMR-effect*). In terms of brain activity, we added the cognitive composite score as a covariate of interest to a *t*-test between the conditions. Model estimations from each participant were taken into two second-level paired samples *t*-tests (CMR > AR and AR > CMR, respectively) with the continuous composite score of cognitive abilities as a covariate of interest.

Moreover, we also controlled for differences in response times and performance, respectively, in relation to the effect of CMR/AR in two separate control analyses. We added the difference in response times between the two conditions, and the difference in performance between the two conditions, respectively as covariates of no interest to two different *t*-tests between the conditions. Model estimations from each participant were taken into a second-level paired samples *t*-test (CMR > AR/AR > CMR) with the difference in response times or performance, respectively, as a covariate of no interest.

## 3. Results

### 3.1 Behavioral Results

#### 3.1.1 Test performance

One week after learning, participants performed significantly better on test items that had been previously learned through CMR compared to AR (see Fig. 2a): *M*_AR_ = .44, *SD* = .21 vs. *M*_CMR_ = .49, *SD* = .24, *t*(71) = −2.06, *p* = .04, 95% CI [−.105, −.002]. In addition, significantly shorter response times were observed for test items learned through the CMR condition compared to AR: *M*_AR_ = 40.9 s, *SD* = 9.4 s vs. *M*_CMR_ = 36.0 s, *SD* = 8.8 s; *t*(70) = 6.44, *p* < .001, 95% CI [3.38, 6.41]. Comparing the subjective post-hoc difficulty ratings averaged over the 12 AR test items and the 12 CMR test items, the difference was not significant: *M*_AR_ = 4.04, *SD* = .92 vs. *M*_CMR_ = 3.93, *SD* = .88; *t*(71) = 1.18, *p* = .24, 95% CI [−.08, .30].

**Figure 2.**
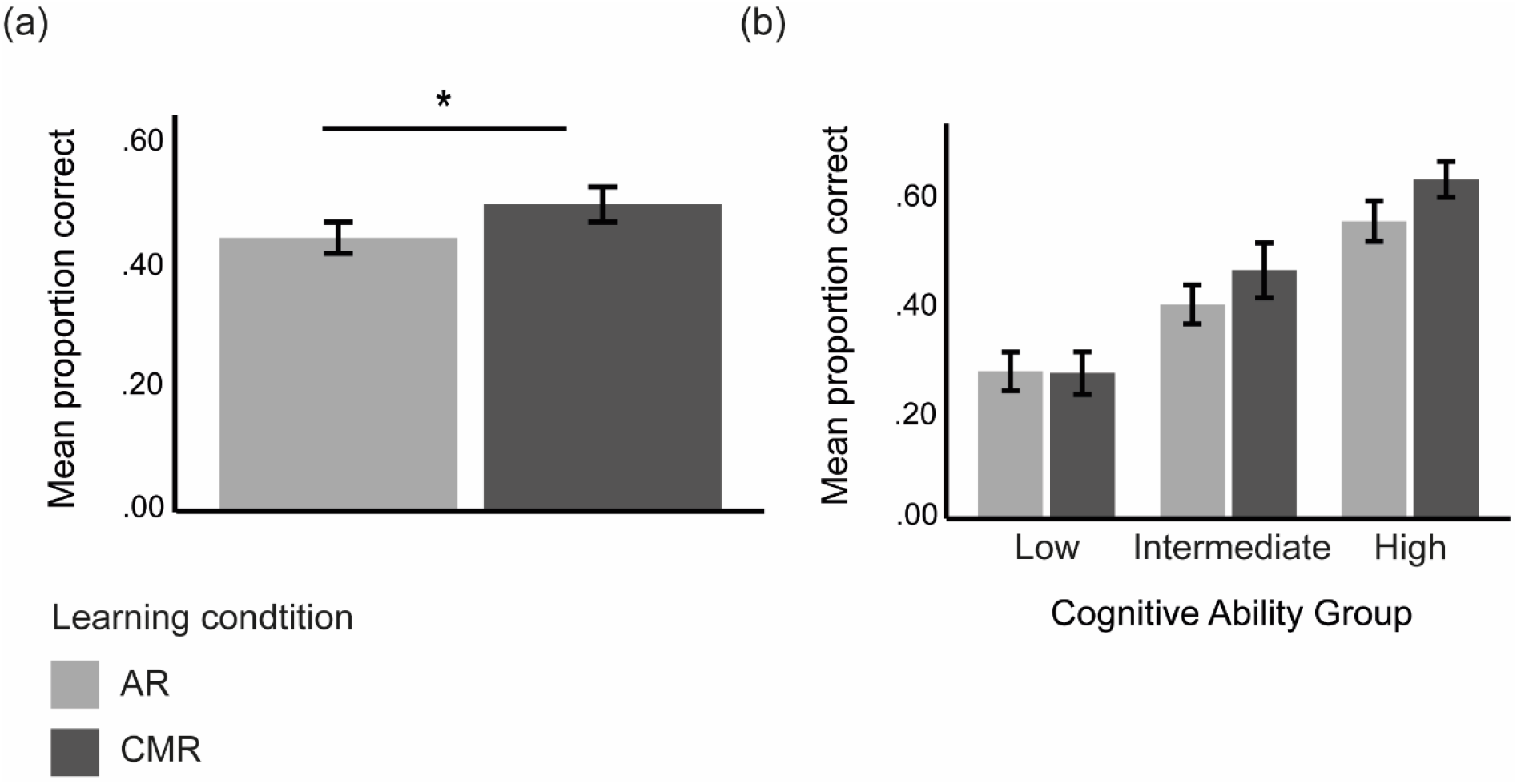
Behavioral performance (proportion correct) during the fMRI test. a) Mean proportion correct on tasks previously learnt with AR and CMR, respectively, b) mean proportion correct on tasks previously learnt with AR and CMR, separately for the three cognitive ability groups (low, intermediate, high). AR = algorithmic reasoning, CMR = creative mathematical reasoning. Error bars: +/- 1 SE. * denotes significant difference at p< .05

To investigate to what extent the benefits on later test performance after practicing with CMR compared to AR depends on cognitive ability, a 2 × 3 mixed model ANOVA was performed. When cognitive ability group (low, intermediate, high) was entered as a between-subjects factor and type of test item (i.e., from tasks that were learned through AR and CMR, respectively) as a within-subjects factor (Fig. 2b) the effect of condition was non-significant, albeit in the same direction as the t-test, CMR > AR (*F*(1, 67) = 3.14, *p* = .08, η_p_^2^= .05). Moreover, we observed a significant main effect of cognitive ability group (low/intermediate/high: *F*(2, 67) = 25.2, *p* < .001, η_p_^2^ = .43). The higher cognitive ability, the better participants performed on the mathematical test independent of CMR/AR (see Fig. 2b). Post-hoc tests revealed significant differences in performance between all three cognitive ability groups (all *p*’s < .023). There was no significant interaction between cognitive ability group and condition (*F*(2, 67) = .81, *p* = .45, η_p_^2^ = .02).

For completeness, we ran a regression analysis to investigate to what extent the continuous cognitive composite score predicted the difference in performance between CMR and AR (performance CMR minus AR: the *CMR-effect*). The continuous cognitive composite score did not significantly predict the CMR-effect, *β* = .15, *t*(69) = 1.24, *p* =.22.

### 3.2 Imaging Results

Next, we turned to analyzing potential effects of learning condition (AR/CMR) and cognitive ability group on brain activation one week after learning. We observed significant main effects of condition (AR/CMR) on brain activity (see Fig. 3 and Table 1 for statistics), but no interaction between condition and cognitive ability group and no main effect of cognitive ability group at the predefined statistical threshold. Main effects of condition (AR/CMR) were mainly observed in a number of fronto-parietal clusters (see Fig. 3 and Table 1). For instance, we observed higher activity in left IFG [−40, 46, 10] and left AG [−42, −60, 48] when participants solved test items previously learned through CMR compared to AR (see Fig. 3a and 3b, and Table 1).

**Figure 3.**
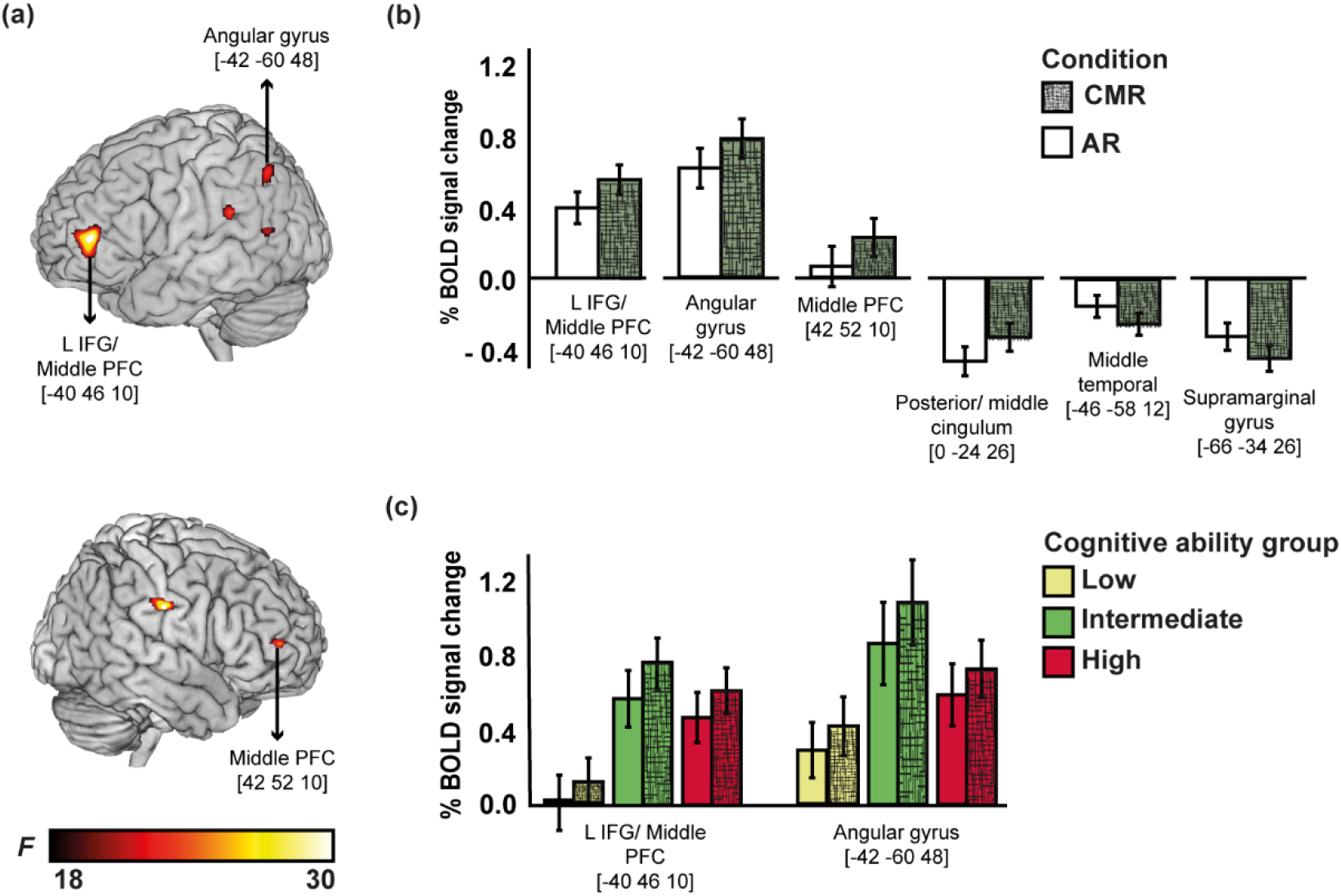
BOLD signal change results from the mixed model ANOVA with learning condition (AR/CMR) as within subject factor and cognitive ability group (low/intermediate/high) as between subject factor. a) Lateral regions exhibiting a main effect of learning condition (see also Table 1); b) Bars illustrating the BOLD signal change when solving test items previously learnt with AR and CMR, respectively. AR = algorithmic reasoning, CMR = creative mathematical reasoning; c) Bars illustrating BOLD signal change in left IFG/mid PFC, and left AG on tasks previously learnt with AR and CMR, separately for the three cognitive ability groups (low, intermediate, high). Patterned bars represent the CMR condition. Error bars: +/- 1 SE.

**Table 1.**
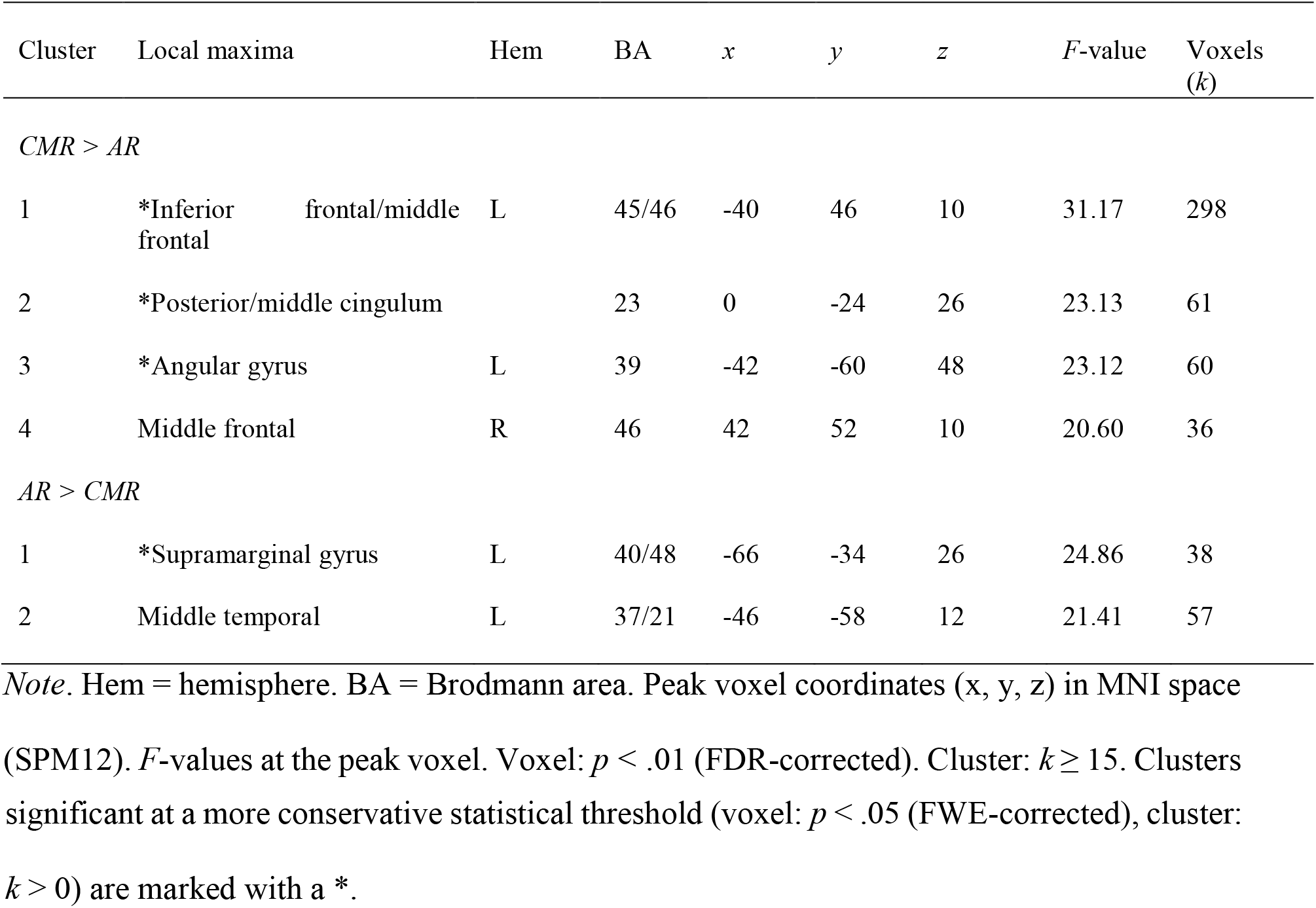
Local maxima for main effect of learning condition (AR/CMR) from the mixed model ANOVA with learning condition (AR/CMR) as within subject factor and cognitive ability group (low/intermediate/high) as between subject factor. AR = algorithmic reasoning. CMR = creative mathematical reasoning.

In addition, significant main effects of learning condition were also seen in right mid PFC, posterior/middle cingulum, supramarginal gyrus and middle temporal cortex, where the latter two clusters were more activated for test items previously learned through AR compared to CMR (Figure 3b, Table 1). Because activation in left AG and left IFG is particularly consistent with results from previous brain imaging studies on mathematics, and for illustrative purposes, signal change in those regions was plotted as a function of cognitive ability group (see Fig. 3c). The same pattern was seen for all three cognitive ability groups, with a tendency for numerically less activation for the lower cognitive ability group compared to the others.

For completeness, entering the continuous cognitive composite score as a covariate to the *t*-test between the conditions (CMR vs AR and vice versa), did not affect the conclusions above. Left IFG/mid PFC, left AG, right mid PFC and posterior/middle cingulum were still more strongly activated when participants solved test items previously learned through CMR compared to AR. Importantly, no clusters covaried with the continuous cognitive composite score at the predefined statistical threshold. The effects observed between CMR and AR remained even when entering differences in performance and reaction time, respectively, as covariates.

## 4. Discussion

Here we hypothesized that constructing your own solutions to mathematical problems with *creative mathematical reasoning* (CMR-tasks; see e.g., Lithner, 2008; Lithner, 2017) should be more beneficial for learning than tasks promoting imitating a provided algorithmic solution template (AR-tasks) because of the initial effort and focus on task-relevant information exerted during learning with CMR. We expected differential recruitment of the left AG region and potentially other fronto-parietal brain regions associated with mathematical cognition (Seghier, 2013; Wirebring, Lithner, et al., 2015; Zhou et al., 2018) when comparing tasks learned through CMR compared to AR one week prior to fMRI scanning. We included cognitive abilitiy group as a factor (Grabner et al., 2007; Hoffman, 2018; Kim, 2020; Zhou et al., 2018) in order to outline whether cognitive abilities interact with the effects of learning method.

With pupils participating in a within-subject intervention in the classroom, followed by a test situation during brain imaging with fMRI one week later, we observed better performance on test items learnt through CMR compared to AR. We thus replicate several previous studies, with a similar task design, in demonstrating a CMR-effect: that CMR-tasks are more effective in fostering mathematical learning than AR-tasks (see e.g., Jonsson et al., 2020a; Jonsson et al., 2016; Jonsson et al., 2014; Norqvist, 2018; Norqvist et al., 2019; Wirebring, Lithner, et al., 2015). The format of the AR-tasks in the experiment, with an introduction, a solution method and a question, is common in school (Jäder, Lithner, & Sidenvall, 2019). The format of the CMR-tasks, without a solution method, is less common, even though problem solving without given solution methods has been proposed in many countries curricula policies as one of the main ways to learn mathematics well (Boesen et al., 2014).

Interestingly, we did not observe an interaction between cognitive ability group and learning method on performance but instead a main effect of cognitive abilities. Individuals with relatively higher cognitive abilities performed better than individuals with relatively lower cognitive abilities, one week after learning. Individual differences in cognitive abilities have been shown to predict academic achievement in various settings (Alloway & Alloway, 2010; Gathercole, Pickering, Knight, & Stegmann, 2004; Kyttälä & Lehto, 2008; Wiklund-Hörnqvist et al., 2016) as well as performance on the type of tasks used in this particular study (Jonsson et al., 2020a; Jonsson et al., 2014; Wirebring, Lithner, et al., 2015).

Brain imaging data makes it possible to observe neural evidence for differences in learning outcomes despite relatively small behavioral differences, especially in relation to the complexity of mathematical thinking (Sohn et al., 2004). In the present study, the brain imaging contrasts differ only in terms of previous training history (i.e. learning condition one week prior to the fMRI test session). We observed significant differences in brain activity in several brain regions, including left IFG and left AG on tasks previously learned through CMR compared to AR. Activity differences in these regions are consistent with other studies on various forms of arithmetical and mathematical thinking (Dehaene et al., 2003; Wirebring, Lithner, et al., 2015; Zamarian et al., 2009; Zhou et al., 2018). Notably, these effects remained when controlling for performance as well as response times. This implies that the brain activity differences mirror processing or representational differences between the two types of tasks (CMR vs AR) rather than being driven by performance differences *per se*.

One interpretation of these observed activation differences is that training with the presumably more effortful CMR-tasks increases the potential to mobilize relevant fronto-parietal control networks at test (cf. Bjork & Bjork, 2011: see Kim, 2020; Wirebring et al. 2015; Zamarian et al. 2009; Zhou et al 2018). However, participants did not perceive the CMR-tasks at test to be more difficult, neither did they spend more time solving CMR test tasks compared to AR test tasks.

Another interpretation concerns enhanced allocation of attention to task-relevant memory during CMR compared to AR test tasks (Cabeza, Ciaramelli, Olson, & Moscovitch, 2008). Activity in the more dorsal part of left AG, that we observe here (*z* = 48), has been suggested to be linked with bottom-up processes during search from semantic memory (Price, 2010; Seghier, 2013) and automatic allocation of attention to memory (Cabeza et al., 2008; Ciaramelli, Grady, & Moscovitch, 2008). Related to this, recent research on mathematical processing suggests that one role of AG in arithmetical processing might be to mediate and control whether or not processes related to the task are employed by targeting relevant networks (Bloechle et al., 2016; Klein, Willmes, Bieck, Bloechle, & Moeller, 2019). Specifically, Bloechle et al. (2016) and Klein et al. (2019) suggests that the left AG act as a ‘circuit breaker’ that adjusts and adapts relative attentional demands in the networks associated with magnitude manipulation and fact retrieval.

Yet a third interpretation of the observed activity differences is related to semantic memory. Left lateralized brain activity in especially IFG and AG is identified as key for semantic memory processing and conceptual representations (Binder & Desai, 2011; Gray, Fry, & Montaldi, 2020; Martin & Chao, 2001). Even though mathematical reasoning is likely to recruit brain regions unique to mathematical content (Amalric & Dehaene, 2016, 2019), left AG has been found to be co-activated with lateral PFC regions, linked to semantic processing of the specific mathematical content (Amalric & Dehaene, 2016). IFG has been suggested as part of a ‘semantic working memory’ system (Martin & Chao, 2001), and evidence exists that higher activity in the left IFG is related to access of available semantic representations (Salami et al., 2010). Hence, learning with CMR might benefit learning and memory by prompting task relevant semantic processes by reactivating integrated mathematical representations related to terminologies and principles.

Potentially mirroring the marked performance differences between the cognitive ability groups, we observed a tendency that individuals with relatively higher cognitive abilities had higher brain activity in left IFG and left AG than individuals with relatively lower cognitive abilities (Fig. 3c), but importantly, there was no main effect of cognitive ability group on brain activation data. This stands in contrast to previous studies identifying a relation between individual differences in cognitive abilities and brain activation in parietal and frontal cortices during mathematical processing (Grabner et al., 2007; Grabner et al., 2011; Gullick et al., 2011; Price et al., 2013; Wirebring, Lithner, et al., 2015). For example, Grabner et al. (2007) found a positive relation between an individual’s mathematical and numerical IQ and signal change in AG during arithmetic problem solving. A key feature distinguishing our study and some of the previous attempts is the focus on non-routine mathematical reasoning (see Fig. S2 for examples of task details) instead of a direct relationship between cognitive abilities and basic arithmetical processing (Grabner et al., 2007; Grabner et al., 2011; Gullick et al., 2011; Price et al., 2013). In our study, the lack of interaction between cognitive ability group and learning condition, both concerning performance and brain activity, suggest that the three cognitive ability groups approach the test tasks with similar neurocognitive strategies. This resonates with previous studies that have failed to observe interactions on brain activity between numerical IQ and problem complexity (Grabner et al., 2007). Even though intervention studies on mathematical learning methods, with ecologically valid didactical tasks, using fMRI at posttest are rare (e.g., Wirebring, Lithner, et al., 2015; see Foisy, Matejko, Ansari, and Masson 2020 for examples and a discussion). Our results open up the possibility that encouraging pupils to engage in more active constructive processes when learning mathematical reasoning might benefit all pupils, even those with relatively lower cognitive abilities (Jonsson et al., 2020a, b). In order to even out performance differences between lower and higher ability pupils it might be important to boost cognitive processing among the lower ability pupils by extending the CMR intervention. Hence, if there is a dose-response relationship between intervention and learning outcome, the lower cognitive ability group could have gained even more in terms of performance and brain activity if the intervention had been longer in duration. Research from related fields is promising in this respect. Donnelly, Huber, and Yeatman (2019) showed that for so-called “struggling” readers there was a clear dose-response relationship, with a linear increase in reading ability as a function of reading intervention intensity.

The majority of brain imaging studies on mathematical cognition to date have focused on relatively simple arithmetical processing (for overviews see e.g. Arsalidou, Pawliw-Levac, Sadeghi, & Pascual-Leone, 2018; Arsalidou & Taylor, 2011; Dehaene et al., 2003; Peters & De Smedt, 2018; Zamarian et al., 2009). Evidence in relation to how brain function and structure subserve the forms of more advanced or non-routine mathematics targeted here remains scarce (e.g., Wirebring, Lithner, et al., 2015; Zhou et al., 2018). The exact localization and direction of the left AG activation reported here differs from the previous study aiming to capture mathematical reasoning processes with CMR and AR (Wirebring, Lithner, et al., 2015). This difference might be the result of a number of factors. For example, while Wirebring, Lithner, et al. (2015) had a between-subject design including only nine different task sets and an average of 23 minutes spent on the learning intervention, the present study had a within-subject design including 12 different task sets and 48 minutes training time per condition (resulting in 96 minutes in total and 24 task sets per participant). The difference with respect to time spent on the learning intervention is considerable. Moreover, the fMRI test session in the present study subjected each individual to 24 test items from the two different learning conditions which might have had a direct effect on left AG activation given the recent notion of AG as an attentional “circuit breaker” balancing retrieval (Bloechle, et al., 2019; Klein et al., 2019) and attentional (Cabeza et al., 2008) demands. Another important factor to consider is that the previous study included a cognitive-perceptual baseline task, where the task was to look at a similar display as the mathematical task display, albeit the task was to judge whether there was a spelling mistake (yes/no). Given the known importance of left AG for processes related to reading and comprehension (e.g. Price, 2010) subtracting this baseline activity from the mathematical task activity (Wirebring, Lithner, et al., 2015) might have canceled out memory related AG activity in unintended ways.

Future research should consider designs allowing for multivariate pattern analysis of brain imaging data to create the possibility to detect more subtle activity differences after learning with CMR and AR in relation to individual differences in cognitive abilities (see e.g., Wirebring, Wiklund-Hörnqvist, et al., 2015). Potentially, processes and representations are less precise one week after learning AR compared to CMR. Moreover, targeting the learning phase is vital in order to investigate a hypothetical gradual build-up of memory representations and process as a function of learning with CMR tasks (e.g., Stillesjo, Nyberg, & Wirebring, 2019). That would enable us to connect to the literature observing an important role for hippocampus for mathematical learning (e.g., Klein et al., 2019; Supekar et al., 2013). Our results suggest that mathematical learning could have a lot in common with concept learning.

## 5. Conclusions

In the present study we demonstrated that constructive mathematical reasoning (CMR) compared to imitative learning procedures (AR) lead to better performance and differential brain activity one week after learning. We observed differences in brain activity in a number of regions including left IFG and left AG after CMR compared to AR and these effects did not interact with cognitive ability. We conclude that encouraging pupils to engage in more active constructive processes when learning mathematical reasoning might have beneficial effects on learning and memory as it might support mobilization of relevant control networks, allocation of attention to task-relevant memory and/or reactivation of integrated mathematical representations related to terminologies and principles.

## Author contributions

BJ, LN, JL, CG & CWH designed research, CG, BJ, CWH performed research, LKW & MA analyzed the data, LKW wrote the first draft of the manuscript, LKW, CWH, SS, JL, CG, MA, LN & BJ wrote sections of the manuscript.

## Data availability statement

The data that support the findings of this study will be made available upon request conditional on approval from the requesting researchers local ethics committee and compliance with the European Union General Data Protection Regulations/all relevant guidelines. The completion of a data transfer agreement signed by an institutional official will be required.

## Acknowledgments

This work was supported by the Swedish Research Council (grant number 2014-2099) and by Umeå School of Education (CWH). The authors wish to thank Tony Qwillbard for programming the data collection platform used during the intervention, Mikael Stiernstedt for support during the data collection, and Mathias Norqvist and the members of UFBI for valuable feedback and support throughout the project.

## Appendix: Supplementary methods

**Figure S1.**
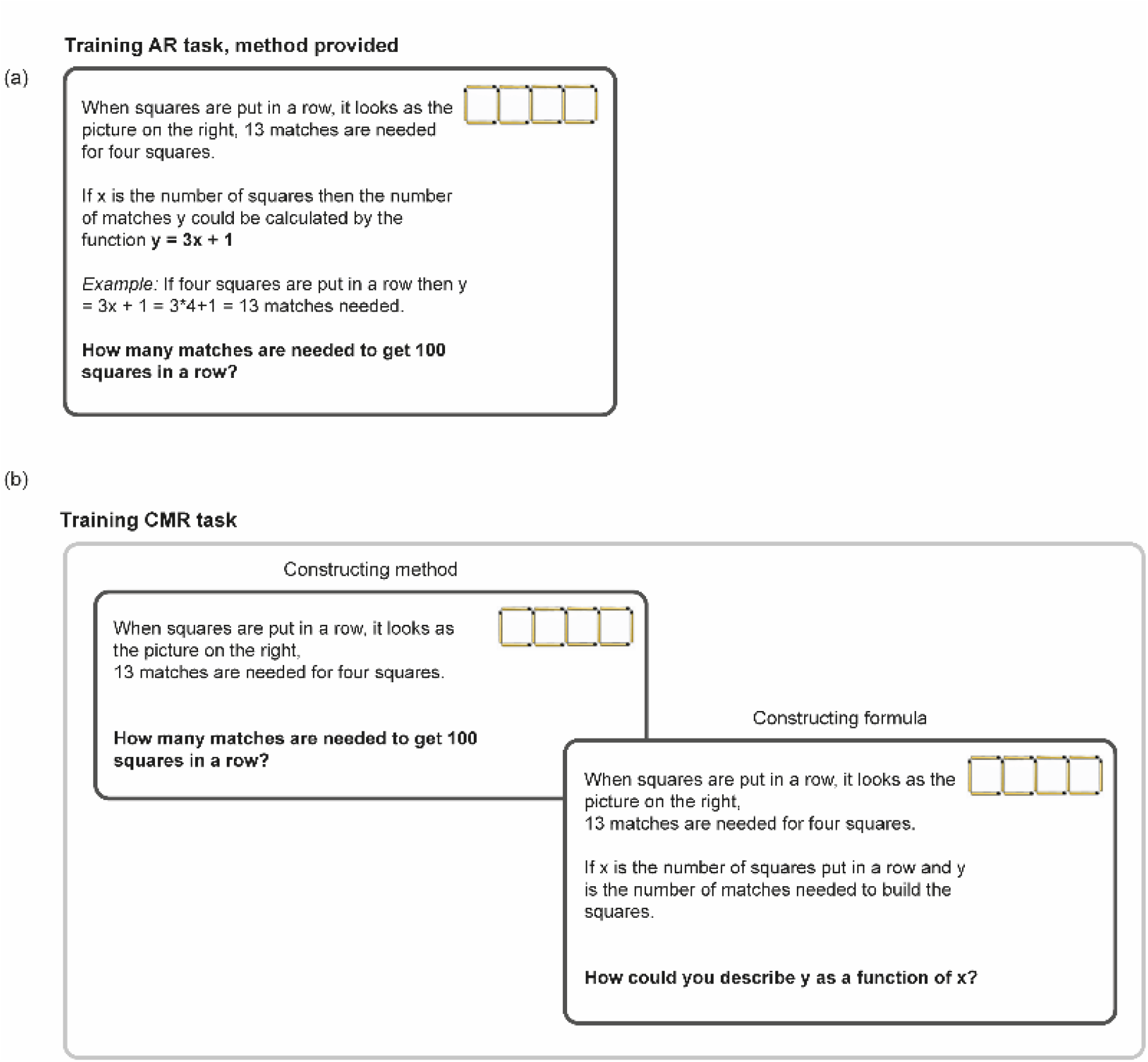
Example of a mathematical task used during the learning intervention. a) In the AR condition, participants were presented with the task (in this case the matches task) as well as a solution template, i.e., a formula and a concrete example how to execute the formula on the specific task. b) In the CMR condition, on the other hand, participants were presented with the task, but did not receive a solution template, i.e., no information regarding the formula nor an example. Instead, on every third trial in a CMR task set participants were given a formula question, asking participants to construct an appropriate formula for the task.

**Figure S2.**
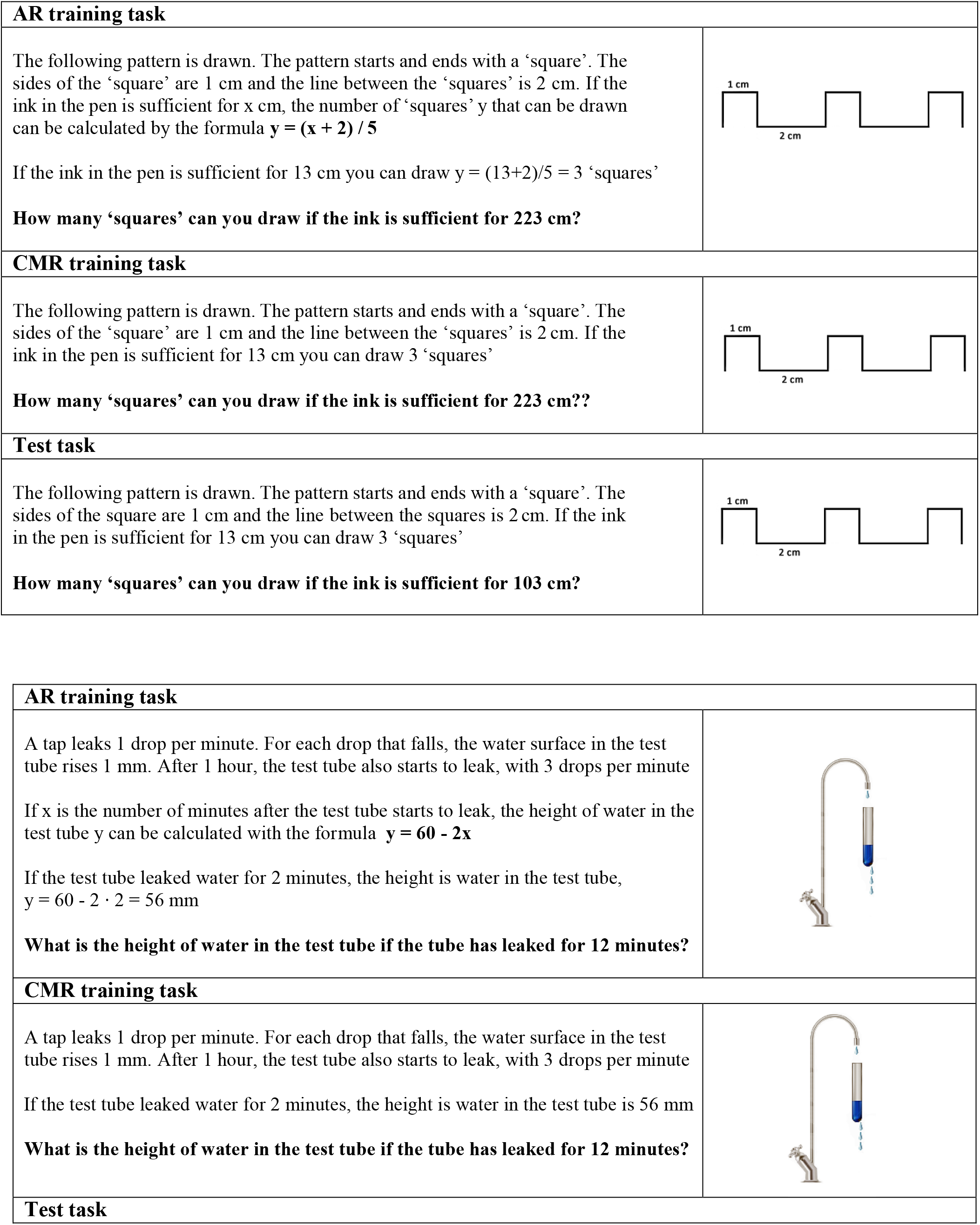
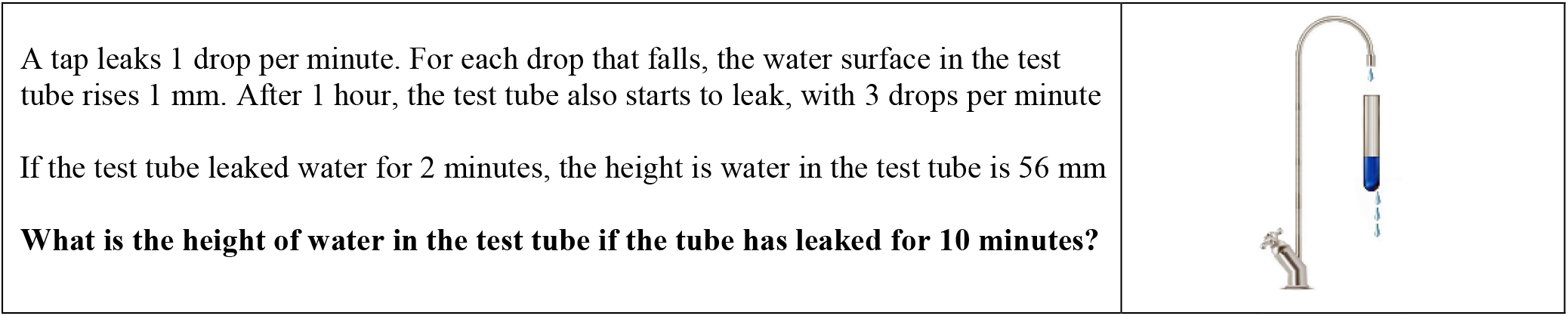
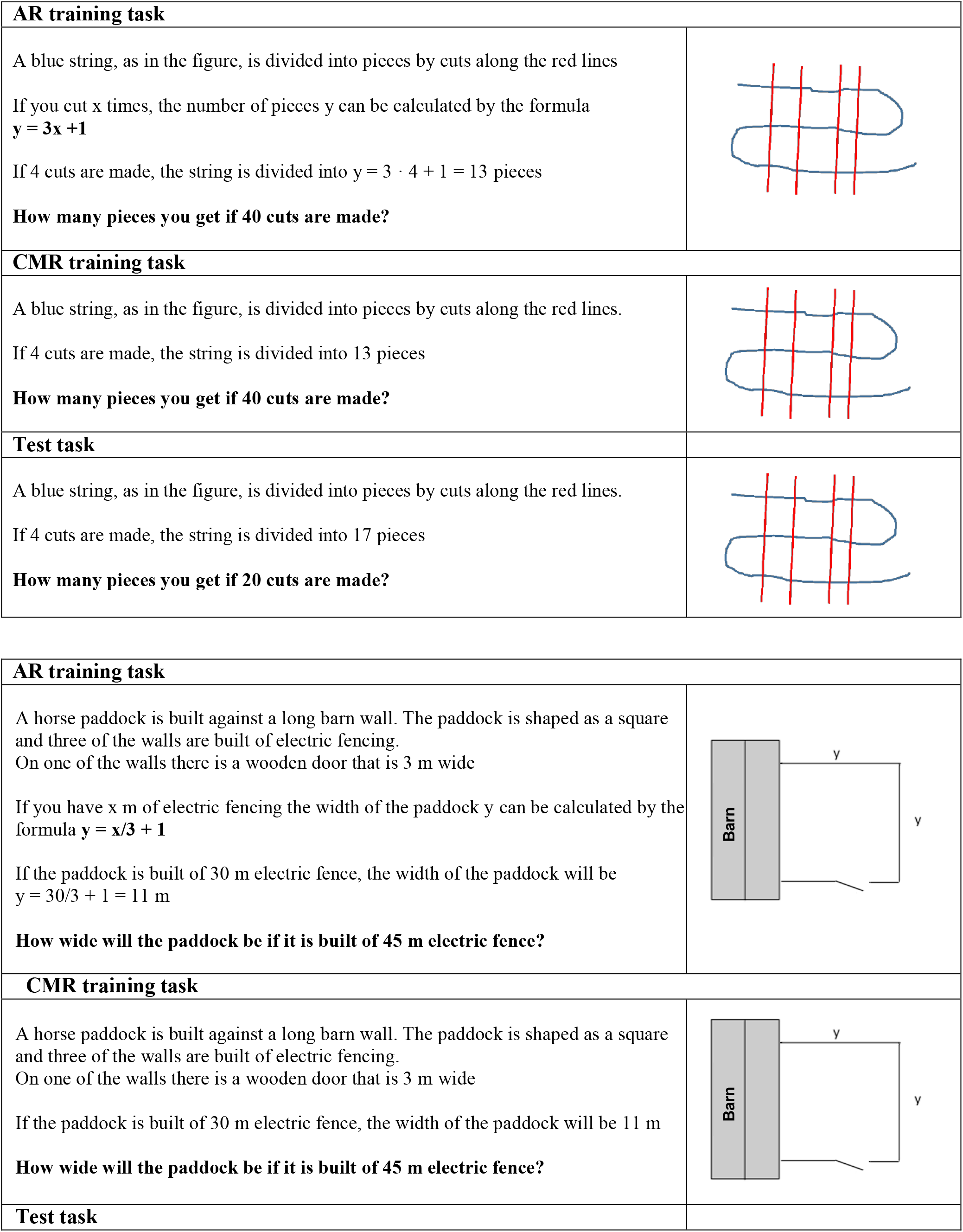
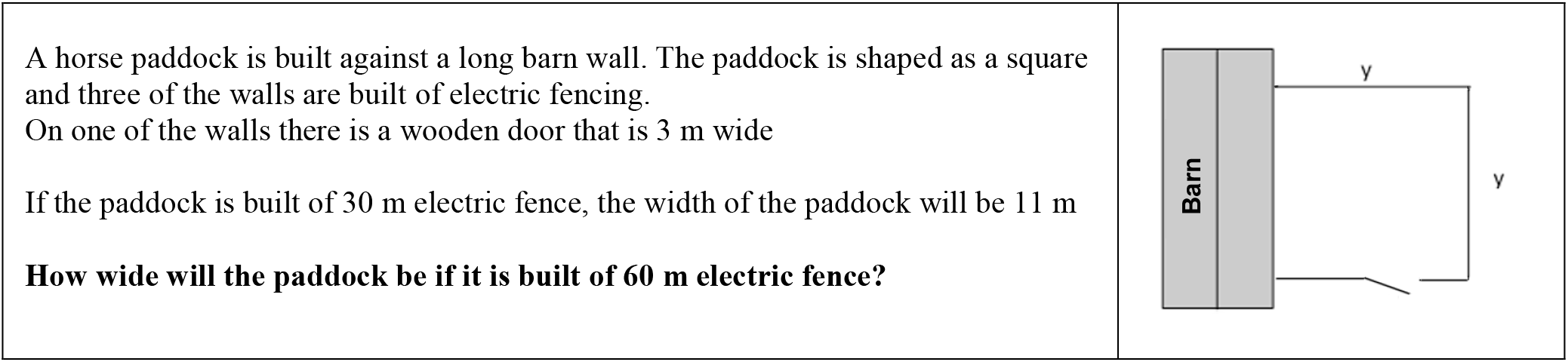
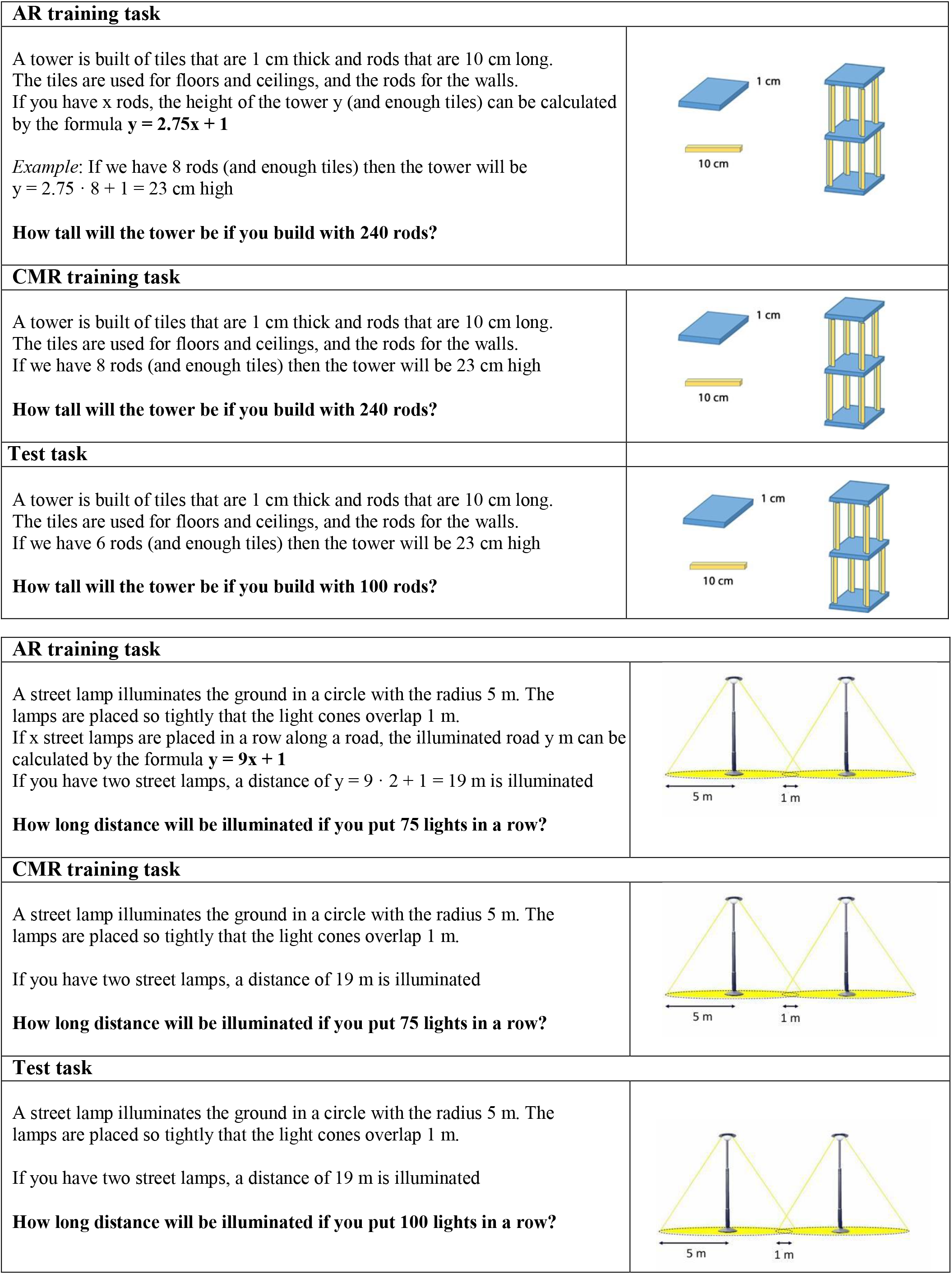
Six additional examples of mathematical tasks used during the learning intervention.

